# Parallel descending neurons differentially encode target and looming information in the hoverfly visual system

**DOI:** 10.64898/2026.11.04.764150

**Authors:** Sarah Nicholas, Karin Nordström

**Affiliations:** Flinders Health and Medical Research Institute, Flinders University, GPO Box 2100, Adelaide, SA, 5001, Australia

**Keywords:** Target motion, descending neuron, target selective descending neuron (TSDN), looming-sensitive descending neuron, insect vision, sensorimotor processing

## Abstract

Detecting small, fast-moving objects is crucial for behaviors such as pursuit, territorial defence, and predator avoidance. At long range, however, potential mates, rivals, and predators produce remarkably similar visual signals, potentially making them difficult to distinguish. Nevertheless, many flying insects, including the hoverfly *Eristalis tenax,* can respond rapidly and appropriately to these ambiguous visual cues, implying specialised neural pathways that extract behaviorally relevant information. Descending neurons link visual processing to motor output, thus transforming visual signals into behavioral actions. In *E. tenax,* both target selective descending neurons (TSDNs) and looming-sensitive descending neurons respond to small targets, yet their respective contributions to target processing remain unclear. Using electrophysiological recordings, we compared receptive field organization and sensitivity to target size and motion direction across these neuronal populations. Although TSDNs and looming-sensitive descending neurons differed systematically in their target coding, a subset of looming-sensitive descending neurons closely resembled TSDNs, sharing small receptive fields and sharp target tuning despite retaining sensitivity to looming stimuli. These findings reveal previously unrecognized diversity among descending neurons tuned to small targets and suggest that parallel descending pathways encode distinct information from similar visual cues. Such an organization may enable common visual inputs to support both target pursuit and the detection of approaching threats.

## Introduction

Responding to small fast-moving objects underpins a diverse range of insect behaviors. In the hoverfly *Eristalis tenax*, for example, looming objects can evoke escape responses^1^, while small targets may trigger male territorial pursuits^2^. Notably, both responses are initiated when objects subtend a small visual angle, yet they can drive opposing behavioral outcomes. At this early stage, visual information about object size is inherently ambiguous, highlighting the importance of precise small target detection as a common sensory signal upon which divergent behavioral responses are built. Understanding how visual information about small objects is represented within descending pathways is therefore central to explaining how insects generate appropriate responses to behaviorally relevant stimuli.

Small target detection is the earliest visual cue to both pursuit and escape responses. Indeed, many animals have highly specialized neurons for this purpose, such as the extensively studied small target motion detectors (STMDs), found in e.g. hoverflies^3–5^ and dragonflies^6–9^. In these species, STMD response profiles are consistent with the size, contrast, and velocity statistics of naturalistic small targets, while exhibiting little or no sensitivity to larger objects or widefield motion^3,6^, making them well suited for pursuit behaviors^10^. Many STMDs also possess small receptive fields suited to precise target localization, while others integrate over larger regions of the visual field^3,4^, suggesting that target information may be represented in parallel even within specialized pathways.

At the level of descending neurons, which link visual processing in the brain to motor output in the thoracic ganglion, both target selective descending neurons (TSDNs)^11,12^ and neurons classically described as looming-sensitive^13–15^, have been shown to respond to translational object motion. TSDNs are strongly selective for small translating targets and show response properties consistent with target-directed behaviors such as pursuit^16,17^. In contrast, looming neurons, here defined as neurons that respond stronger to looming stimuli than to motion-less controls, are classically linked to collision detection and escape behaviors, exemplified by the extensively studied LGMD/DCMD pathway^18–20^. In this framework, responses to small objects and translating trajectories are generally interpreted in the context of monitoring object motion, trajectory and collision risk rather than pursuit and interception^14,21^. Nevertheless, during the final stages of pursuit, time to contact is likely required to initiate capture, and may be derived through looming cues^22^. Indeed, a subset of TSDNs characterized in dragonflies respond to looming-like features^11,23^. Together, these observations suggest that sensitivity to small targets and looming stimuli may not represent mutually exclusive functional specializations, but rather complementary components of parallel pathways responding at different stages of an encounter and supporting behaviors such as pursuit, capture and escape.

While the target encoding properties of TSDNs are increasingly well characterized^11,12,16,17,23–25^, providing insight into their likely role in target pursuit, the target response properties of looming-sensitive descending neurons remain comparatively understudied, despite evidence that some respond robustly to small targets^13,14,26^. Consequently, it remains unclear whether target information is represented similarly across these pathways, or whether variation in target sensitivity within and between neuronal classes reflects distinct functional roles.

Here, we address this question by directly comparing target processing in two classes of descending neurons in the hoverfly *Eristalis tenax*, TSDNs and looming-sensitive descending neurons. Throughout this study, we restrict the term TSDN to neurons that respond selectively to small translating targets with minimal sensitivity to larger objects, looming stimuli or widefield motion. Using published and unpublished electrophysiological recordings, we analyze receptive field structure and tuning for target size and direction to assess population-level similarities and differences between these neuron classes. By explicitly comparing how target information is represented across descending pathways, our study reveals that a subset of looming-sensitive descending neurons share key target response properties with the well-characterized TSDNs, whilst the remaining looming-sensitive descending neurons exhibit considerable diversity. These findings suggest that target and looming cues are integrated differently across descending pathways and may ultimately support distinct behavioral functions.

## Results

### Differential sensitivity to looming stimuli separates two classes of target tuned descending neurons

In this study, we compiled a dataset of 233 descending neurons from 220 previously published recordings^13,17,25–27^ and 13 new extracellular recordings. We assessed each neuron’s responses to small moving targets and looming stimuli and used stringent classification criteria to define neuronal populations for subsequent analyses. Neurons exhibiting little or no response to looming stimuli (*l/|v|* = 10 ms, peak response <30 spikes/s; red data, Figures 1A, 1C and 1D) were classified as TSDNs, whereas neurons exhibiting strong looming responses (peak response >100 spikes/s; blue data, Figures 1B, 1C and 1D) were classified as looming-sensitive descending neurons. Neurons with intermediate responses (30–100 spikes/s; black data, Figures 1C and 1D) were excluded from further analyses. Consistent with previous definitions^13^, neurons here identified as looming-sensitive exhibited substantially stronger responses to the looming stimuli than to a motion-less appearance control (Figure S1). In addition to responding strongly to looming stimuli, many looming-sensitive descending neurons also exhibited robust responses to small targets. However, across the dataset, target-evoked responses were significantly weaker in looming-sensitive descending neurons than in TSDNs (Figure 1E).

**Figure 1.**
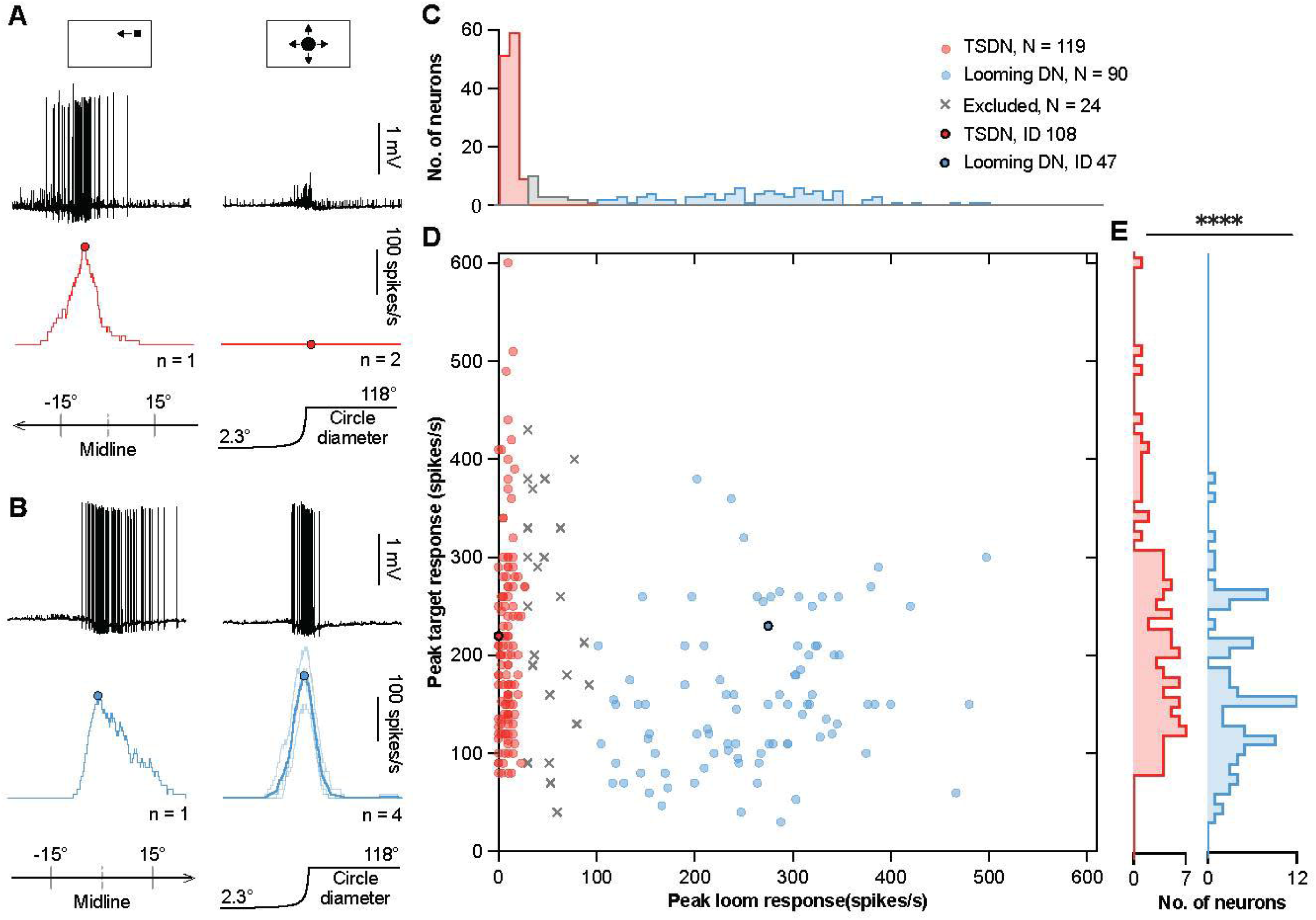
Target responsive descending neurons can be separated by their responses to looming stimuli. (A) Representative responses of a TSDN (neuron ID 108) to a small moving target (left) and a looming stimulus (right). Top, raw electrophysiology recordings (scale bar, 1 mV); middle, corresponding spike histogram, with thin lines representing individual repeats (*n*), thick lines showing the mean, and filled circles indicating the peak response (scale bar, 100 spikes s⁻¹); bottom left, target trajectory; bottom right, loom diameter. (B) Representative responses of a looming-sensitive descending neuron (Looming DN; neuron ID 47) as in panel A. (C) Distribution of peak looming responses for all recorded neurons. Neurons were classified as TSDNs (red, peak looming response <30 spikes s⁻¹; *N* = 119) or looming DNs (blue, peak looming response >100 spikes s⁻¹; *N* = 90). Neurons with intermediate looming responses were excluded from further analyses (gray, N = 24). (D) Peak target responses plotted against peak looming responses for all neurons. The example neurons shown in (A) and (B) are indicated with black outlines. (E) Distributions of peak target responses for TSDNs and looming DNs. TSDNs exhibited significantly greater target responses than looming DNs (*p* < 0.0001, Mann-Whitney U test). See also Figure S1 and S2.

### Looming-sensitive descending neurons have larger, more lateral and more ventral receptive fields

We next examined the spatial organization of target sensitive receptive fields in TSDNs and looming-sensitive descending neurons. Using methods previously applied to TSDNs^17^, we quantified target receptive field properties for all neurons included in the analysis (Figure S2). These measurements characterize only target responsive receptive fields, as looming-sensitive descending neurons also possess distinct receptive fields for looming and widefield stimuli^26^. Individual TSDNs were typically centered near the visual midline and exhibited relatively narrow horizontal extents (Figure 2A), whereas looming-sensitive descending neurons generally had broader receptive fields whose centers were distributed further laterally (Figure 2B). This was reflected in significantly greater receptive field widths for looming-sensitive descending neurons (blue) than for TSDNs (red; Figure 2C). Receptive field centers of looming-sensitive descending neurons were also significantly more eccentric (Figure 2D), resulting in reduced overlap with the visual midline compared to TSDNs (Figure 2E).

**Figure 2.**
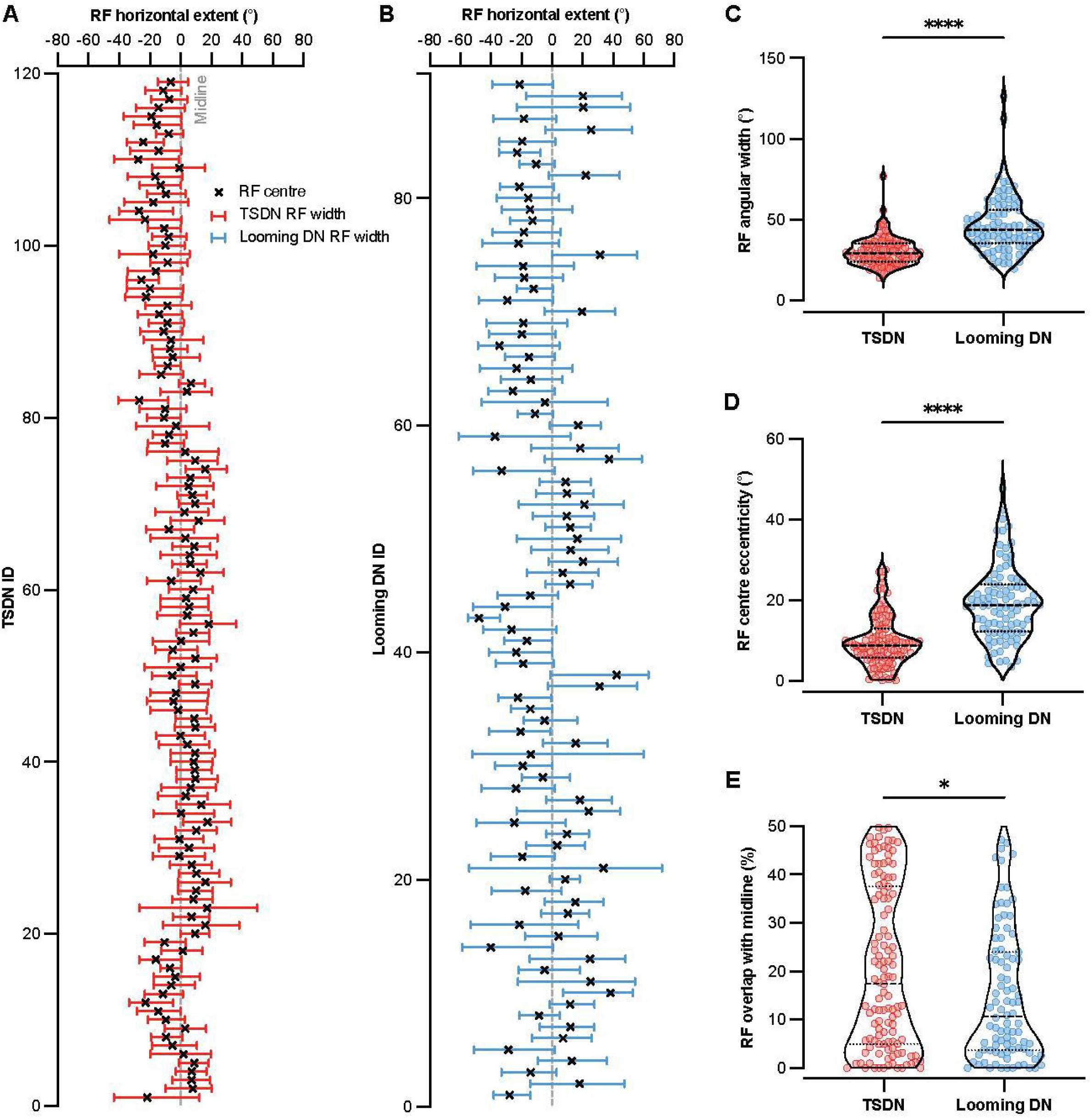
Receptive fields of looming-sensitive descending neurons are wider and more lateral than those of TSDNs. (A and B) Horizontal receptive field (RF) extent of individual TSDNs (A) and looming-sensitive descending neurons (looming DNs; B). Black crosses indicate RF centers, colored bars indicate RF width, and the dashed line indicates the visual midline. (C) Distribution of RF angular widths for TSDNs and looming DNs. Looming DNs had significantly wider RFs than TSDNs (*p* < 0.0001, Mann-Whitney U test). (D) Distribution of RF center eccentricities, calculated as the absolute angular distance between the RF center and the visual midline. Looming DNs had significantly more lateral RF centers than TSDNs (*p* < 0.0001, Mann-Whitney U test). (E) Distribution of RF overlap with the visual midline. Looming DNs exhibited less RF overlap with the midline than TSDNs (*p* < 0.05, Mann-Whitney U test). In all panels, TSDNs are shown in red and looming DNs in blue. Violin plots show the distribution of the data, with central lines indicating the median and the first and third quartiles.

Similarly, the vertical extent of receptive fields also differed. Individual TSDNs were centered dorsal to the visual equator and exhibited relatively narrow vertical extents (Figure 3A), whereas looming-sensitive descending neurons exhibited receptive field centers both above and below the visual equator and had broader vertical extents (Figure 3B). This was reflected in significantly greater receptive field heights for looming-sensitive descending neurons (blue) than for TSDNs (red; Figure 3C). Receptive field centers of looming-sensitive descending neurons were also located at lower elevations (Figure 3D), resulting in greater overlap with the visual equator compared to TSDNs (Figure 3E). While these differences were significant at the population level, the distributions overlapped considerably, with some looming-sensitive descending neurons exhibiting receptive fields similar in size and position to those of typical TSDNs (e.g. compare Figure S2G with Figures S2F and S2I).

**Figure 3.**
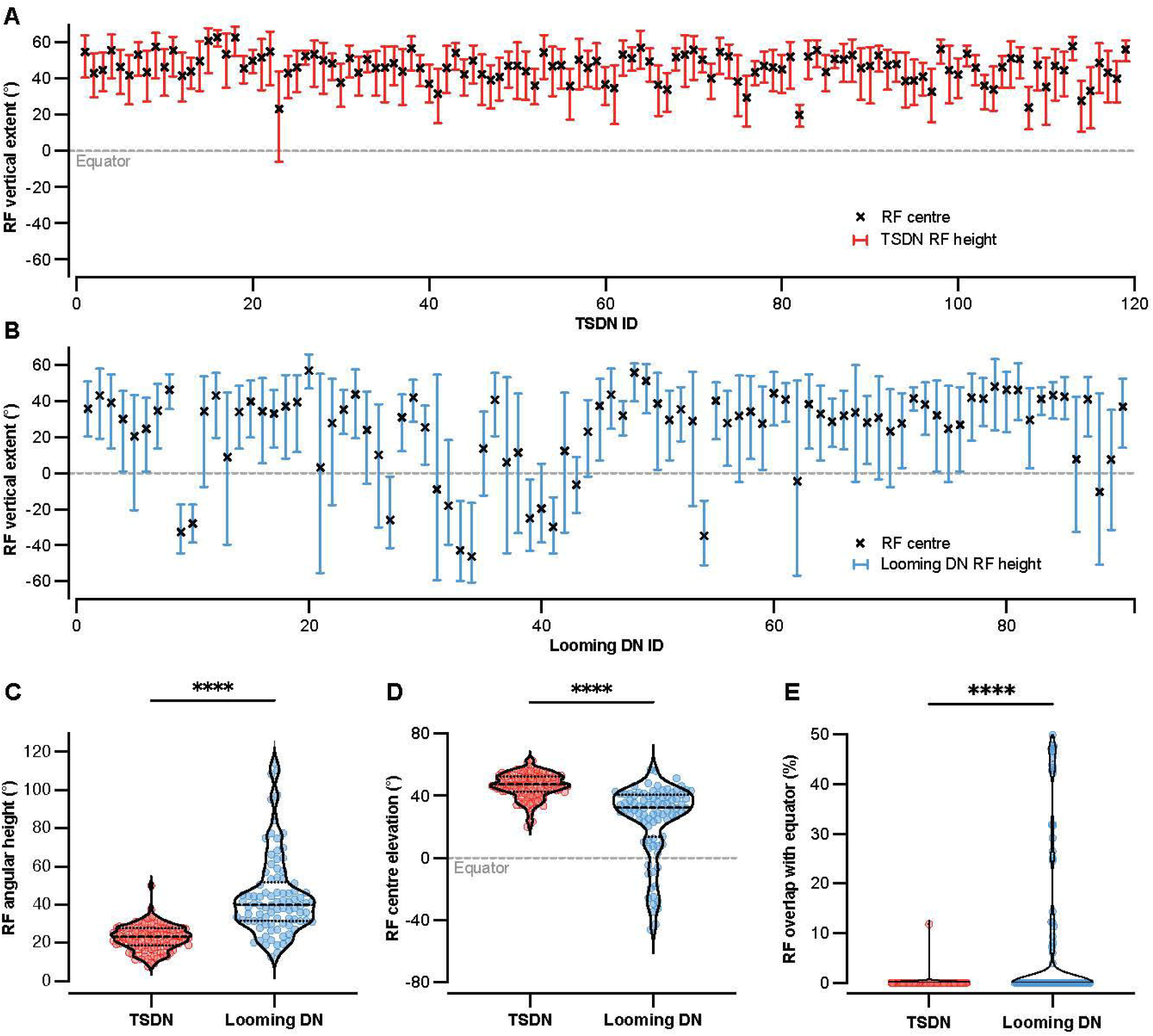
Receptive fields of looming-sensitive descending neurons are taller and more ventral than those of TSDNs. (A and B) Vertical receptive field (RF) extent of individual TSDNs (A) and looming-sensitive descending neurons (looming DNs; B). Black crosses indicate RF centers, colored bars indicate RF height, and the dashed line indicates the visual equator. (C) Distribution of RF angular heights for TSDNs and looming DNs. Looming DNs exhibited significantly taller RFs than TSDNs (*p* < 0.0001, Mann-Whitney U test). (D) Distribution of RF center elevations, calculated as the angular distance between the RF center and the visual equator. Looming DNs had significantly more ventral RF centers than TSDNs (*p* < 0.0001, Mann-Whitney U test). (E) Distribution of RF overlap with the visual equator. Looming DNs exhibited more RF overlap with the equator than TSDNs (*p* < 0.0001, Mann-Whitney U test). In all panels, TSDNs are shown in red and looming DNs in blue. Violin plots show the distribution of the data, with central lines indicating the median and the first and third quartiles.

### Looming-sensitive descending neurons exhibit reduced directionality and increased angular variance

We next investigated whether differences in receptive field organization were accompanied by differences in directional sensitivity, including overall directionality and spatial variation in preferred direction across the receptive field. Spatial variation in preferred direction was quantified using angular variance (V, Figure S2D), calculated from the distribution of local preferred directions measured at locations with local motion sensitivity exceeding 50% of its maximum value (red lines, Figure S2D). Angular variance quantifies the consistency of local preferred directions across the receptive field, with higher values indicating greater spatial variability. The strength of directional tuning was quantified using a directionality index (DI, Figure S2E), calculated from a single directional tuning curve (red line, Fig S2E) derived from responses at the same locations.

Looming-sensitive descending neurons exhibited significantly lower directionality and higher angular variance indices than TSDNs, indicating both less selective and less spatially consistent directional tuning (Figure 4A, 4B). These differences in directionality and angular variance were accompanied by differences in the distribution of preferred directions (PD, Fig. S2D) between the two populations. Preferred direction was calculated as the circular mean of local preferred directions measured at locations with local motion sensitivity exceeding 50% of its maximum value (red lines, Figure S2D). A greater proportion of TSDNs exhibited a horizontal preferred direction (85%) compared with looming-sensitive descending neurons (52%), whereas vertical preferred directions were more common among looming-sensitive descending neurons (37%) than TSDNs (14%) (Figure 4C–4H). The remaining neurons were classified as non-directional (11% of looming-sensitive descending neurons and 1% of TSDNs) due to either high angular variance (V > 0.7) or a low directionality index (DI < 0.3; Figure 4A–4F). Among neurons with a vertical preferred direction (orange and brown, S3A–S3D), only looming-sensitive descending neurons possess receptive field centers below the equator (Fig. S3C, D), all of which preferred upward motion (brown, Figure S3D). In contrast, neurons with center positions above the equator showed no consistent preference for upward or downward motion (orange and brown, Figure S3A-S3D). TSDNs with a horizontal preferred direction largely preferred motion away from the visual midline based on receptive field center position, a trend that was less apparent in looming-sensitive descending neurons (purple and green, Figure S3E-S3H).

**Figure 4.**
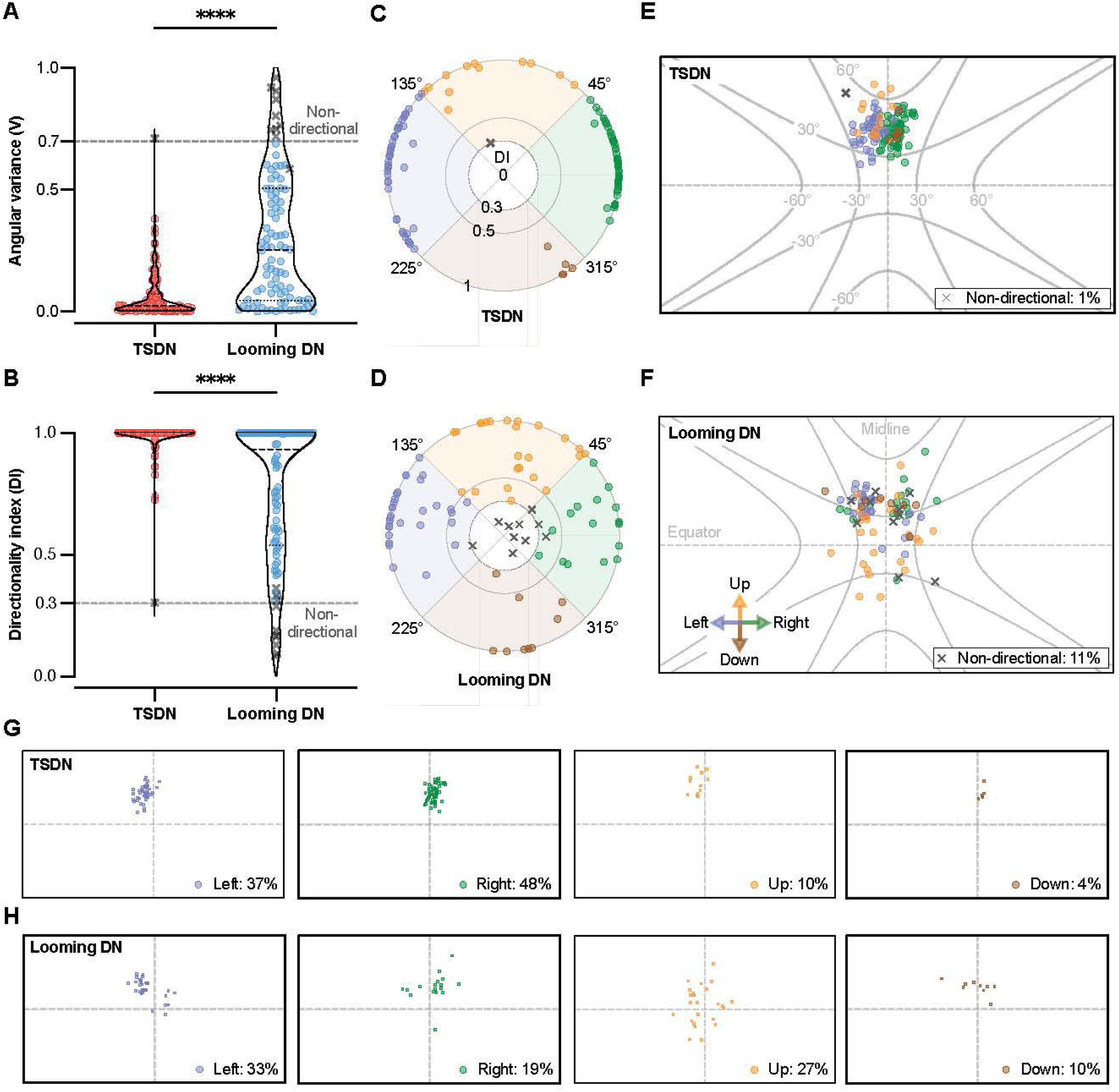
Looming-sensitive descending neurons exhibit increased angular variance and reduced direction selectivity. (A) Distribution of receptive field angular variance (V) for TSDNs and looming-sensitive descending neurons (Looming DNs). Looming DNs (blue) exhibited significantly greater angular variance than TSDNs (red, *p* < 0.0001, Mann-Whitney U test). The dashed line indicates the threshold used to classify neurons as non-directional. (B) Distribution of directionality index (DI) values for TSDNs and looming DNs. Larger values indicate stronger directionality. Looming DNs exhibited significantly lower DI values than TSDNs (*p* < 0.0001, Mann-Whitney U test). The dashed line indicates the threshold used to classify neurons as non-directional. (C and D) Polar plots showing the preferred direction of individual TSDNs (C) and looming DNs (D). Symbols indicate individual neurons, color-coded according to preferred direction. Radial distance represents directionality index (DI; shown in B), and gray crosses indicate neurons classified as non-directional according to the thresholds shown in (A) and (B). (E and F) Location of receptive field centers for TSDNs (E) and looming DNs (F), with symbols indicating individual neurons, color-coded according to preferred direction. Gray crosses indicate non-directional neurons, with the percentage in each grouping shown. (G and H) Receptive field centers of individual TSDNs (G) and looming DNs (H), grouped according to preferred direction. Percentages indicate the proportion of directional neurons preferring leftward (purple), rightward (green), upward (yellow), or downward motion (brown). See also Figures S2 and S3.

### A subset of looming-sensitive descending neurons exhibit TSDN-like characteristics

Although looming-sensitive descending neurons and TSDNs exhibit significant differences in target receptive field characteristics at the population level (Figure 2–4), considerable overlap is observed across all measured metrics. To quantify the extent of this overlap, we used principal component analysis (PCA) to define a TSDN feature space and applied a squared Mahalanobis distance (D²) threshold to classify neurons relative to the TSDN distribution. Lower D² values indicate receptive field properties most similar to those of TSDNs, whereas higher values indicate progressively greater divergence from the TSDN distribution. Most TSDNs clustered at low D² values, with 91.6% of TSDNs falling below a threshold of D² = 16.92 and fewer than 1% exhibiting D² values >100 (red, Figure 5A). In contrast, approximately 44% of looming-sensitive descending neurons exhibited D² values >100, although 24.4% fell below the D² threshold (blue, Figure 5A).

**Figure 5.**
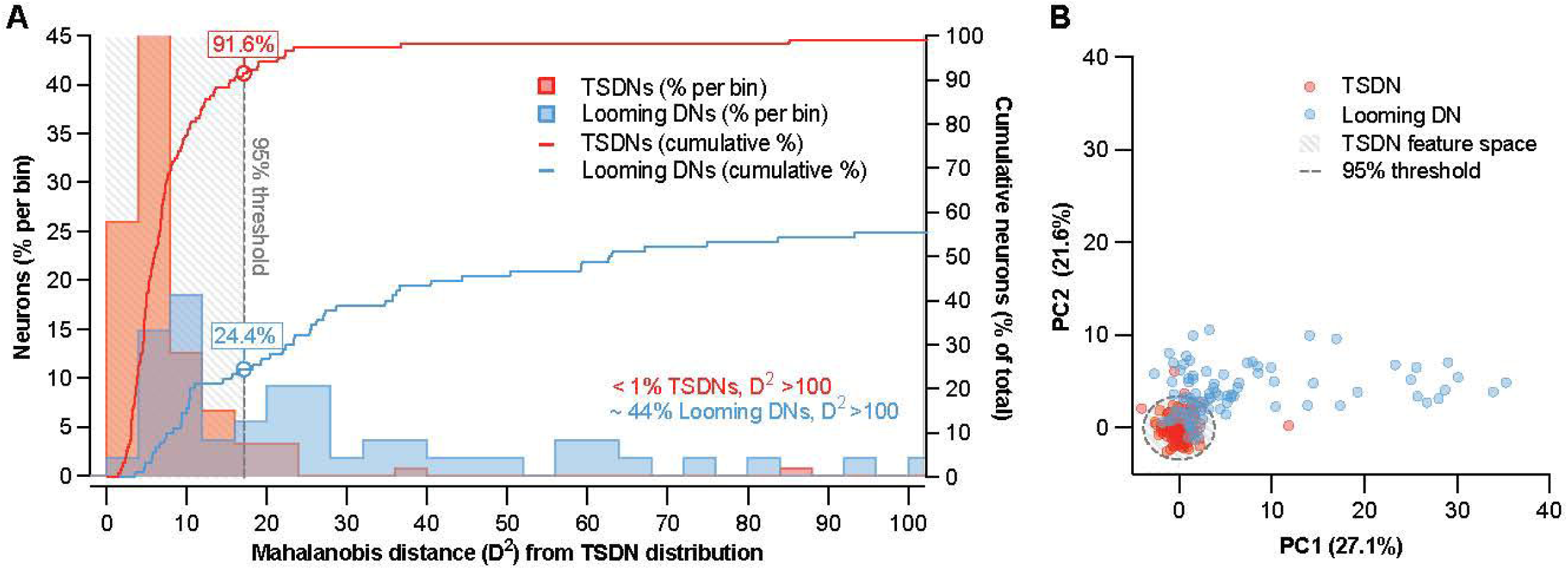
Looming-sensitive descending neurons partially overlap with the TSDN feature space. (A) Distribution of Mahalanobis distances (D²) from the TSDN distribution for TSDNs (red) and looming-sensitive descending neurons (blue, Looming DNs). Histograms show the percentage of neurons within each D² bin, while lines show cumulative percentages. The cross-hatched region and dashed line indicate the 95% confidence boundary of the TSDN feature space (D² = 16.92). Approximately 92% of TSDNs and 24% of looming DNs fell within this boundary. (B) Principal component analysis (PCA) of neuronal feature space. Each point represents a single neuron, plotted according to the first two principal components (PC1, 27% variance explained; PC2, 22% variance explained). The cross-hatched region and dashed contour indicate the TSDN feature space. While many looming DNs (blue) fell outside the TSDN feature space, a subset overlapped with the TSDN cluster (red).

These results indicate that while some looming-sensitive descending neurons exhibit TSDN-like target receptive field properties, many others differ substantially from the TSDN distribution. Consistent with this, projection into PCA space revealed that a subset of looming-sensitive descending neurons occupied the same feature space as TSDNs, whereas the remaining neurons extended well beyond the TSDN cluster (Figure 5B). This enabled further subdivision of looming-sensitive descending neurons into two groups: “TSDN-like” looming-sensitive descending neurons (low D², D² < 16.92) and “non-TSDN-like” looming-sensitive descending neurons (high D², D² > 16.92).

“TSDN-like” looming-sensitive descending neurons are less responsive to looming stimuli

Having identified a subset of looming-sensitive descending neurons with receptive field properties similar to TSDNs (gray dashed area, Figure 5A), we next investigated whether these neurons exhibit additional TSDN-like response characteristics and whether they differ from “non-TSDN-like” looming-sensitive descending neurons. TSDNs (red), “TSDN-like” looming-sensitive descending neurons (gray) and “non-TSDN-like” looming-sensitive descending neurons (blue) responded similarly to small bars, with firing rates increasing as bar height increased and reaching an initial peak at bar heights of just a few degrees (Figure S4). However, whereas TSDN responses declined sharply for larger bars and remained low (red, Figure S4), looming-sensitive descending neurons exhibited a secondary increase in responsiveness at large bar heights (blue and gray, Figure S4). This secondary rise was evident in both “TSDN-like” and “non-TSDN-like” looming-sensitive descending neurons and was most pronounced at bar heights exceeding 40° (blue and gray, Figure S4).

Consistent with these differences, the bar height at response onset (Figure 6A) and the bar height corresponding to the initial peak response (Figure 6B) did not differ significantly between the groups. However, “non-TSDN-like” looming-sensitive descending neurons exhibited significantly larger response offset bar heights than TSDNs (Figure 6C), indicating that they maintained responses to a broader range of stimulus sizes. In contrast, “TSDN-like” looming-sensitive descending neurons did not differ significantly from TSDNs (Figure 6C).

**Figure 6.**
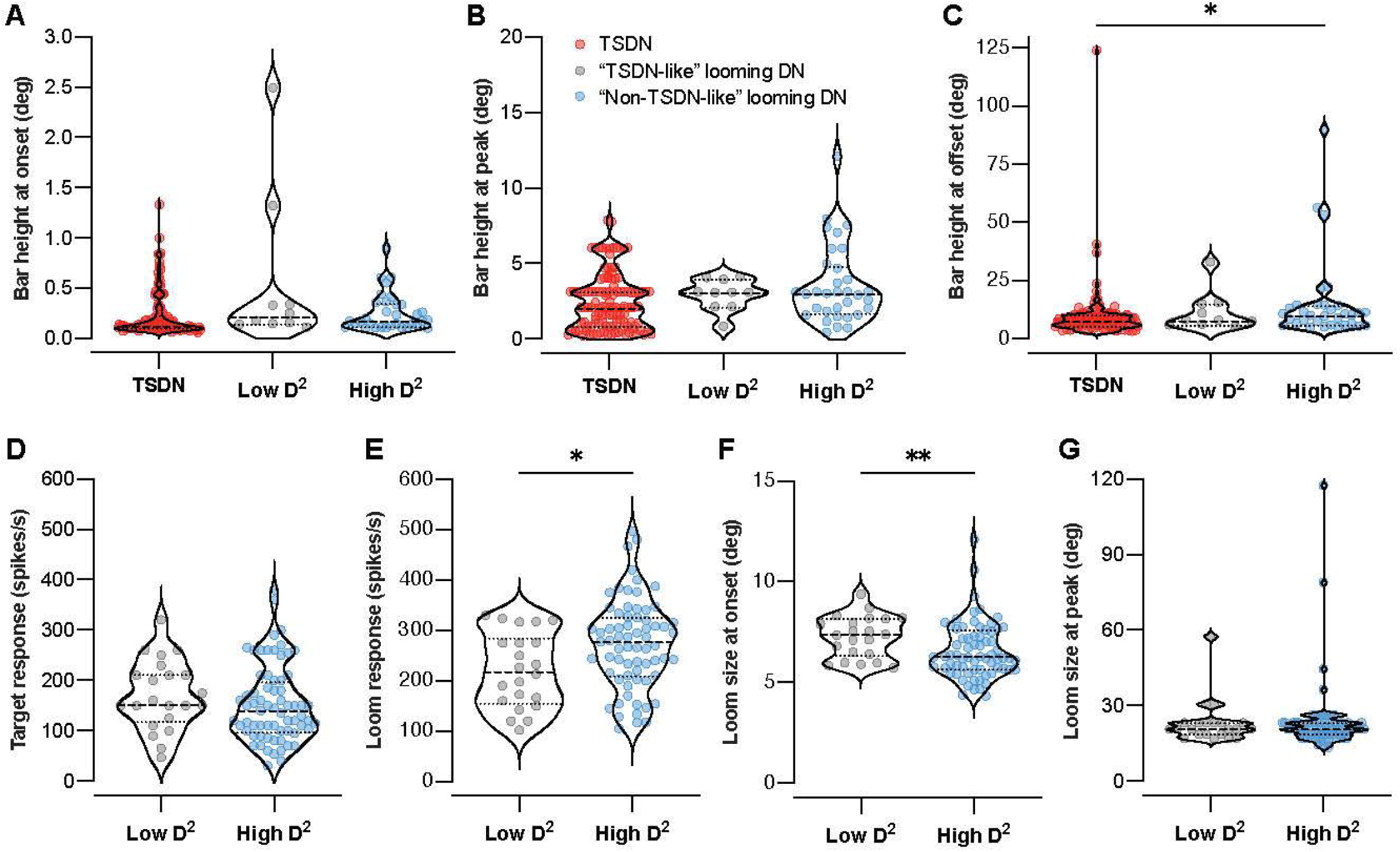
“Non-TSDN-like” looming-sensitive descending neurons respond at smaller loom sizes and exhibit stronger responses to looming stimuli. (A-C) Comparison of bar size tuning between TSDNs (red), “TSDN-like” looming-sensitive descending neurons (“TSDN-like” looming DN; low D²; gray), and “non-TSDN-like” looming-sensitive descending neurons (“non-TSDN-like” looming DN; high D²; blue). (A) Bar height at response onset. (B) Bar height at peak response. (C) Bar height at response offset. Significance was investigated with Kruskal-Wallis tests. (D-G) Target and looming response characteristics of looming-sensitive descending neurons. (D) Peak target response. (E) Peak looming response. “Non-TSDN-like” looming DNs exhibited significantly stronger responses to looming stimuli than “TSDN-like” looming DNs (*p* < 0.05, Mann-Whitney U test). (F) Loom size at response onset. “Non-TSDN-like” looming DNs responded significantly earlier than “TSDN-like” looming DNs (*p* < 0.01, Mann-Whitney U test). (G) Loom size at peak response. In all panels, violin plots show the distribution of the data, with central lines indicating the median and the first and third quartiles. See also Figure S1 and S4.

Next, we compared looming evoked responses between “TSDN-like” and “non-TSDN-like” looming-sensitive descending neurons. Although the two groups exhibited similar peak responses to small moving targets (Figure 6D, the combined data correspond to the target responses of looming-sensitive descending neurons shown in Figure 1), “non-TSDN-like” looming-sensitive descending neurons (blue) had a significantly greater peak response to looming stimuli compared to “TSDN-like” looming-sensitive descending neurons (gray; Figure 6E).

To further explore differences in looming responses, we fitted a spline function to the mean response at each circle size during looming stimulus expansion and used the resulting function to estimate the circle size at response onset and peak response (Figure S1D). “Non-TSDN-like” looming-sensitive descending neurons responded significantly earlier, with response onset occurring at smaller circle sizes, than “TSDN-like” looming-sensitive descending neurons (Figure 6F). However, both groups reached their maximum response at similar circle sizes during the looming stimulus expansion (Figure 6G). Together, these results indicate that “non-TSDN-like” looming-sensitive descending neurons are both more responsive (Figure 6E) and more sensitive (Figure 6F) to looming stimuli than their “TSDN-like” counterparts.

## Discussion

We found that hoverfly descending neurons responsive to small targets can be separated into two populations, TSDNs and looming-sensitive descending neurons, based on their sensitivity to looming stimuli (Figure 1). These populations differ not only in their responses to looming stimuli, but also in the organization of their target receptive fields and their directional tuning properties (Figure 2–4). Despite these population level differences, substantial overlap exists between the two groups, with multivariate analysis identifying a subset of looming-sensitive descending neurons with receptive field characteristics that fall within the TSDN feature space (Figure 5). Although these “TSDN-like” looming-sensitive descending neurons respond similarly to small targets, they exhibit weaker and less sensitive responses to looming stimuli than other looming-sensitive descending neurons (Figure 6). Together, these findings suggest that target tuned descending neurons may be organized into overlapping functional pathways. TSDNs and “TSDN-like” looming-sensitive descending neurons appear specialized for precise spatial localization and tracking of small moving targets, whereas “non-TSDN-like” looming-sensitive descending neurons trade spatial precision for broader target sensitivity and enhanced detection of approaching objects. These pathways may therefore support distinct behavioral functions, while retaining common visual strategies for detecting behaviorally relevant objects.

### Target detection during pursuit

TSDNs are widely considered strong candidate neurons for the control of target pursuit owing to their sharp size tuning, small dorso-frontal receptive fields and strong directional selectivity^16,17^. However, analyses based on reconstructed pursuit trajectories suggest that the response properties of the known TSDN population are insufficient to explain the full range of visual inputs experienced during pursuit^17^. This is particularly evident during the final stages of an approach, when the retinal image of a female conspecific rapidly expands as the distance between the insects closes^28^, potentially exceeding the preferred size range of TSDNs^17^ and transitioning from a small translating target to a looming stimulus. These observations suggest that, while TSDNs play a crucial role in pursuit, additional target responsive descending pathways are likely required to maintain target detection across the full pursuit trajectory.

“TSDN-like” looming-sensitive descending neurons represent a strong candidate for filling this gap. Consistent with their classification as “TSDN-like” neurons, they fall within the TSDN feature space and possess the small dorso-frontal receptive fields, low angular variance and strong directional selectivity characteristic of classical TSDNs (Figure 5), while also retaining the sharp size tuning required for target detection during pursuit (gray, Figure 6A–6C). Unlike TSDNs, however, they additionally respond to looming stimuli. Importantly, responsiveness to translating targets declines at a median angular size of 8.1° (gray, Figure 6C), very close to the median angular size at which looming responses emerge (7.4°; gray, Figure 6F), indicating these neurons remain responsive as the dominant visual cue shifts from small target motion to retinal image expansion. In this way, “TSDN-like” looming-sensitive descending neurons may complement classical TSDNs by supporting target detection across a broader portion of the pursuit trajectory, particularly during the final stages of approach and capture.

### Early warning for approaching objects

While the detection of small targets is essential for pursuit behaviors, sensitivity to small moving objects is likely to support a broader range of visually guided behaviors. Indeed, the presence of STMDs in female hoverflies, which do not exhibit the pursuit behavior typical of males, indicates that small target detection is not uniquely tied to pursuit, but instead serves a broader behavioral function^10^. One such role may be the early detection of approaching objects. Approaching animals initially generate little retinal image expansion and are therefore more readily detected at a distance through their translational motion across the visual field. Consistent with this idea, behavioral studies have shown that escape responses of hoverflies feeding on flowers are only partly explained by looming cues, suggesting that additional visual signals contribute to the detection of approaching threats^1^. The “non-TSDN-like” looming-sensitive descending neurons described here may contribute to such a mechanism. Unlike TSDNs and “TSDN-like” looming DNs, “non-TSDN-like” looming-sensitive descending neurons, lack the small dorso-frontal receptive fields, low angular variance, and strong directional selectivity characteristic of these populations (Figure 5). As a result, they respond to small targets across a larger portion of the visual field and over a broader range of motion directions. Such properties may increase the likelihood of detecting approaching animals at an early stage, albeit with reduced spatial and directional specificity. In addition, “non-TSDN-like” looming-sensitive descending neurons respond more strongly to looming stimuli (blue, Figure 6E) and begin responding when the looming stimulus subtends a smaller angular size (blue, Figure 6F). Together, these properties could provide both early detection of approaching threats and robust signaling as retinal image expansion increases, making these neurons well suited to support rapid escape responses.

### Similar visual cues, opposing behavioral outcomes

The visual cues generated by a distant moving animal are inherently ambiguous, as they may signal either an opportunity for pursuit or a threat requiring avoidance. For example, the behavioral response to an incoming conspecific may differ depending on whether an individual is a female foraging at a flower^1^ or a male engaged in territorial surveillance^2^. The present study suggests that hoverflies may address this challenge through parallel descending pathways that extract distinct information from the same visual scene. While TSDNs and “TSDN-like” looming-sensitive descending neurons appear specialized for small target detection and tracking, “non-TSDN-like” looming-sensitive descending neurons trade spatial precision for broader sensitivity to small moving and approaching objects. Such an organization could allow common visual inputs to support both pursuit and avoidance behaviors, while preserving the distinct sensory information relevant to each task. Rather than encoding behavioral decisions directly, these pathways may provide complementary representations of approaching objects that differ in their balance between precision and sensitivity. The resulting response would then depend on how these parallel signals are integrated by downstream circuits and interpreted within the current behavioral context.

## Resource Availability

## Lead Contact

Requests for further information and resources should be directed to and will be fulfilled by the lead contact, Sarah Nicholas.

## Materials availability

This study did not result in the generation of new material

## Data and code availability

- All data and custom analysis code have been deposited to DataDryad and will be released upon publication.

## Acknowledgements

We thank Biomedical Engineering at SAHLN and the Botanic Gardens of Adelaide for their ongoing support. This research was funded by the US Air Force Office of Scientific Research (AFOSR, FA9550-23-1-0473) and the Australian Research Council (ARC, DP230100006 and DP250100698).

## Author contributions

Conceptualization, S.N. and K.N.; methodology, S.N. and K.N; Investigation, S.N; writing – original draft, S.N.; writing – review & editing, S.N. and K.N; funding acquisition, K.N.; resources, K.N; supervision, K.N.

## STAR Methods

## Experimental model and study participant details

*Eristalis tenax* hoverflies were reared from eggs laid by wild-caught females collected under permit in Adelaide, South Australia, with larvae and adults maintained under standard laboratory conditions as described previously^29^. Recordings were obtained from adult males approximately 3 months of age.

## Method details

### Data collection

We systematically screened and reanalyzed all available extracellular electrophysiological recordings from hoverfly descending neurons previously identified as either target-selective descending neurons (TSDNs) or looming-sensitive descending neurons, using the definitions described by Nicholas *et al.*^13^. Recordings included in this study were obtained across a series of previous investigations from our laboratory^13,17,25–27^ and supplemented with previously unpublished data. Only neurons for which a complete target receptive field was recorded, and looming sensitivity was tested were included (see details below). This yielded a dataset of 233 unique neurons recorded from 215 individual hoverflies, including 220 previously published and 13 new recordings.

Detailed descriptions of animal preparation, electrophysiological recording techniques, and data acquisition are provided in the original publications^13,17,25–27^. In brief, adult hoverflies were immobilized in a dorsal side up orientation, and a small hole was made in the thoracic cuticle to expose the cervical connective. The connective was elevated and stabilized using a small wire hook, and extracellular recordings were made from descending neurons using a sharp tungsten microelectrode (2 MΩ, polyimide-insulated; Microprobes). Neural signals were amplified (×1000), band-pass filtered between 10 and 3000 Hz using a DAM50 differential amplifier (World Precision Instruments), with further noise reduction via a HumBug (Quest Scientific). Signals were digitized using a PowerLab 4/30 interface (ADInstruments) at a sampling rate of 40 kHz. Individual units were isolated offline in LabChart 7 Pro (ADInstruments) using spike sorting based on the amplitude and width of individual action potential waveforms.

### Visual stimuli

Visual stimuli were presented on a linearized 165 Hz LCD monitor (2560 × 1440 pixels; Asus, Taipei, Taiwan) using custom MATLAB software (FlyFly 3.2) incorporating the Psychophysics Toolbox^30,31^. The monitor was positioned perpendicular to the animal at a viewing distance of 6.5 cm, subtending 155° × 138°, with a mean illuminance of 200 lux.

### Small target receptive field mapping

All neurons were initially identified based on their responses to a small, black moving target before undergoing detailed receptive field mapping using methods previously described^3,13^. In brief, a 3.5 × 3.5 mm black square (3 × 3° at screen center) moved across the display in the four cardinal directions at 209 mm s⁻¹ (116° s⁻¹ at screen center) and was presented at 20 evenly spaced azimuths and elevations, with a minimum inter-stimulus interval of 0.5 s. When receptive field mapping was repeated, responses were averaged across trials.

Peak target responses were derived from these mapping datasets. Firing rates were calculated for each trajectory using a 100 ms sliding window spike histogram, and peak target response was defined as the maximum firing rate observed across all stimulus locations and directions (left panels, Figure 1A, 1B).

To characterize target receptive fields, the mean spike rate was calculated for each trajectory across 20 bins (142 ms each), incorporating an assumed neural response latency of 18 ms. We then subtracted the spontaneous firing rate, defined as the firing rate during the 0.48 s preceding stimulus onset. This produced a 20 × 20 response matrix for each direction of motion (Figure S2A). Responses to the four motion directions were then averaged and interpolated to achieve a five-fold increase in spatial resolution of the receptive field map (Figure S2B).

Using MATLAB’s *polyshape* function, we determined the 50% contour line (red line, Figure S2C) and measured receptive field center, width and height from this boundary, with all subsequent angular measurements reported after correcting for display geometry and distortion (gray lines, Figure S2C). When multiple disconnected polyshapes were identified, measurements were obtained only from the largest receptive field region (Figure S2H). Receptive field width and height were measured as the maximum horizontal and vertical extents of the 50% response boundary (black bars, Figure S2F–S2K). Receptive field center coordinates were defined as the centroid of the 50% contour, determined using MATLAB’s *centroid* function (red or blue circles, Figure S2C, S2F–S2K). Receptive field center azimuth and elevation correspond to the angular distances of this centroid from the visual midline and visual equator, respectively. Receptive field eccentricity was calculated as the absolute value of the center azimuth, assuming that neurons recorded from the left and right sides of the brain represent matched bilateral pairs.

Midline overlap was quantified as the percentage of receptive field width extending across the visual midline, while equator overlap was quantified as the percentage of receptive field height extending across the visual equator. Values were set to 0% for receptive fields that did not cross the respective axis. For receptive fields crossing the axis, overlap was calculated as the angular distance from the axis to the nearest receptive field edge, divided by receptive field width or height and expressed as a percentage.

To characterize direction selective properties of receptive fields, responses to each direction of motion at every location within the 20 x 20 receptive field matrix were fit with a cosine function to estimate the local preferred direction (LPD) and local motion sensitivity (LMS, Figure S2D). Locations with LMS values greater than 50% of the maximum LMS for that neuron were included in subsequent analyses. For these locations, LPD and LMS were represented as vectors, with vector angle indicating LPD and vector length indicating LMS (arrows, Figure S2E–S2K). Receptive field preferred direction (PD, Figure S2D) was defined as the circular mean of included LPDs, calculated using *circ_mean* from the CircStat MATLAB toolbox^32^. Angular variance (V, Figure S2D) was used to quantify spatial variability in local preferred direction and was calculated as the circular variance of these same LPDs, using *circ_var* functionCircStat MATLAB toolbox, ^32^, with higher values indicating greater spatial variability.

Whereas LPD and LMS were calculated locally for each position in the receptive field matrix, overall directionality was quantified by averaging responses for each direction of motion across locations with LMS values greater than 50% of the maximum LMS for that neuron. A sinusoidal function was then fit to the resulting tuning curve, and the directionality index (DI, Figure S2E) was calculated as:

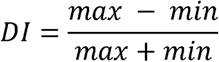

where *max* and *min* denote the maximum and minimum values of the fitted sinusoidal function, respectively. Negative fitted minima were constrained to zero prior to DI calculation, as spike frequency cannot be negative (red line, Figure S2E). To compensate for the ventral side up orientation of each animal during recordings, the receptive fields were rotated to display them with the dorsal visual field up.

### Looming sensitivity

Looming stimuli were generated as previously described^13,20^, using an *l/|v|* of 10 ms. A black disc expanded from 1° to 117° over 1 s and remained at full size for a further 1 s (blue line, Figure S1B), with successive presentations separated by at least 60 s. An appearance control consisted of a stationary black disc (117° diameter) that appeared on the screen and remained visible for 1 s (gray line, Figure S1B). Peak responses for looming (Peak_loom_, blue circle, Figure S1A) and appearance control (Peak_ctrl_, gray circle, Figure S1A) stimuli were defined as the maximum firing rates in trial averaged spike histograms generated using a 100 ms sliding window (right panels, Figure 1A, 1B and Figure S1A). Looming selectivity (LS, Figure S1C) was calculated as:

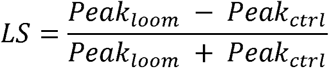

For analyses of response tuning as a function of stimulus size, firing rates were calculated separately for each display frame (6.06ms; 165Hz refresh rate). Each response measurement corresponded to a single stimulus size during the looming stimulus after correcting for an assumed neural response latency of 18 ms. Mean firing rates were then plotted as a function of stimulus size, and a spline fit was used to determine the stimulus size at response onset (10% of peak response) and peak response (Figure S1D).

### Size tuning

Size selectivity was assessed by drifting a black bar through the center of each neuron’s receptive field in its preferred horizontal direction of motion^13,17,26^. The bar travelled across a white background at 209 mm s⁻¹ (116° s⁻¹ at screen center). Bar width was fixed at 3 mm (3° at screen center), while height varied from 0.2 mm to the full height of the display (335 mm). Consecutive stimulus presentations were separated by a minimum interval of 4 s. Bar height in visual degrees was calculated after correcting for display geometry and distortion relative to the stimulus trajectory and the receptive field center of each neuron.

Peak responses to each bar height were defined as the maximum firing rates in trial averaged spike histograms generated using a 100 ms sliding window. Peak firing rates were then plotted against bar height, and a spline fit was used to determine the bar heights at response onset, peak and offset (Figure S4). Response onset was defined as the first bar height at which the spline fit reached 10% of the peak. If the response to the smallest bar tested elicited a response exceeding this threshold, onset was assigned to that bar height (Figure S4C). Response peak was defined as the first local maximum in the spline fit as bar height increased. Response offset was defined as the first bar height at which the spline fit returned to 10% of the peak response after the peak. If the fit did not return to this threshold, offset was assigned to the bar height corresponding to the minimum response following the peak (Figure S4B).

### Population-level analyses

To assess the similarity of looming-sensitive descending neurons to classical TSDNs, we first defined a multidimensional TSDN phenotype space using nine receptive field and response properties: peak target response, angular variance, receptive field center azimuth, receptive field center elevation, directionality index, equator overlap, receptive field height, receptive field width and midline overlap. These variables were normalized using the mean and standard deviation of the TSDN population. Mahalanobis distance (D²) from the TSDN population was then calculated for all neurons. Looming-sensitive descending neurons with D² values falling within the 95% confidence boundary of the TSDN phenotype space, defined using a χ² distribution with nine degrees of freedom (corresponding to the nine variables included in the analysis), were classified as “TSDN-like” (low D²), whereas neurons outside this boundary were classified as “non-TSDN-like” (high D²). Principal component analysis was performed on the normalized TSDN dataset, and looming descending neurons were projected onto the resulting axes for visualization.

### Quantification and statistical analysis

Statistical analyses were performed in Prism 11.0.1 (GraphPad Software). Throughout the text, *n* refers to individual stimulus repetitions, whereas *N* refers to individual neurons. Each neuron was assigned a unique identification number (ID) following sorting by preferred direction. These IDs were maintained throughout the analyses and figures. Unless otherwise stated, data are presented as median and interquartile range (IQR). Violin plots show the distribution of the data; central lines indicate the median and the first and third quartiles. Comparisons between two groups were performed using two-tailed Mann-Whitney *U* tests. Comparisons among three or more groups were performed using Kruskal-Wallis tests. Exact sample sizes, statistical tests, and *p* values are reported in the corresponding figure legends. Significance values are indicated as follows: NS for *p*>0.05, one star (*) for *p*<0.05, two stars (**) for *p*<0.01, three stars (***) for *p*<0.001, four stars (****) for *p*<0.0001.

