## Supplementary Figures for "Parallel descending neurons differentially encode target and looming information in the hoverfly visual system"

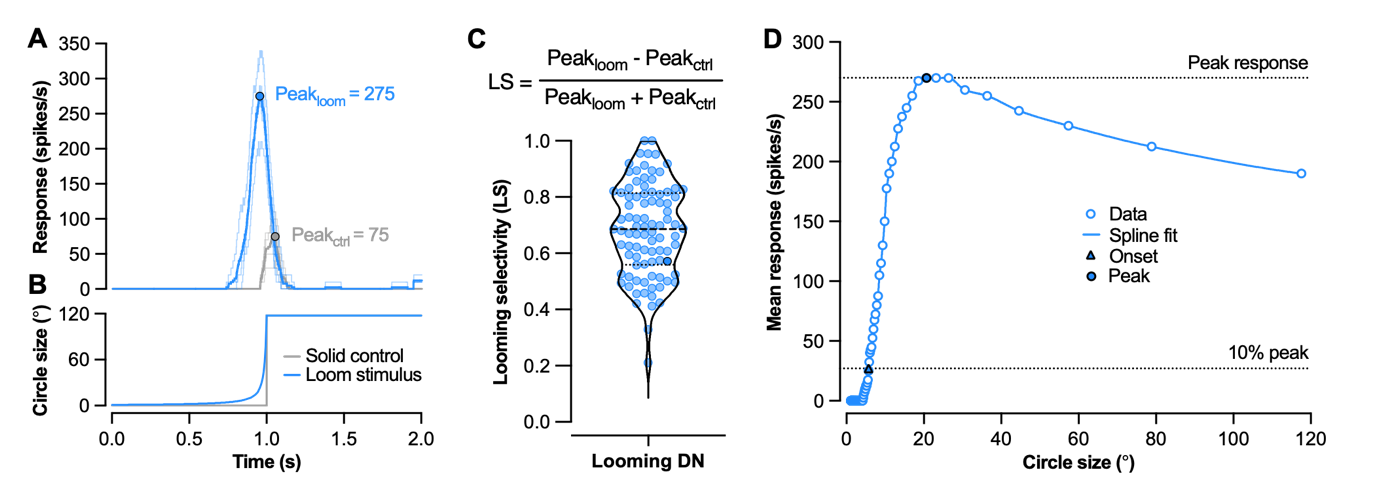
**Figure S1. Quantification of looming selectivity and extraction of loom response parameters.** Supports Figure 1 and 6. (A) Example responses of a looming DN (ID 47; also shown in Figure 1B) to a looming stimulus (blue) and a motion-less control (grey). Histograms show firing rate, with thin lines representing individual trials, the thick line showing the mean response, and filled circles indicating the peak response. (B) Angular size of the looming and motion-less control stimuli shown in (A). (C) Distribution of looming selectivity (LS) across all looming DNs, calculated as the normalized difference between the peak responses to looming and control stimuli, central lines indicating the median and the first and third quartiles. (D) Determination of stimulus angular size at response peak and onset for the example neuron shown in (A). Mean response is plotted as a function of stimulus angular size (open circles). A spline fit (blue line) was used to interpolate responses across stimulus sizes. Peak response size was defined as the angular size at which the fitted response reached its maximum (filled circle). Response onset size was defined as the angular size at which the fitted response first exceeded 10% of the peak response (triangle).

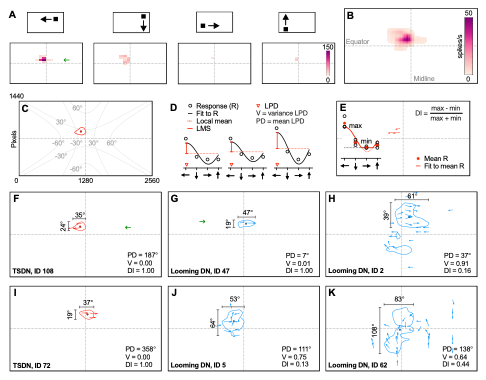
**Figure S2. Receptive field mapping and quantification of receptive field properties.** Supports Figure 1 and 4. (A) Example responses of a TSDN (ID 108) to a small target moving in the four cardinal directions across a 20 × 20 grid of spatial positions spanning azimuth and elevation. Heatmaps show mean spike rate for leftward, downward, rightward, and upward motion at each location within this grid. (B) Receptive field map generated from spatially interpolated mean responses across all motion directions. Dashed lines indicate the visual equator and midline. Color scale denotes firing rate (spikes s⁻¹). (C) The receptive field boundary was defined as the 50% response contour (red) from the interpolated map in (B), with the circle indicating the receptive field center. Grey contours indicate display geometry and associated spatial distortion, which was corrected when calculating angular measurements. (D) Determination of local preferred direction (LPD, red triangle), local motion sensitivity (LMS, red line segment), preferred direction (PD) and angular variance (V). At each spatial position, responses (R) to the four cardinal directions were fitted with a cosine function (black line). LPD was defined as the direction corresponding to the peak of the fitted function, and LMS as the difference between the peak fitted response and the local mean response (red dashed line). The circular mean and variance of LPDs, measured at locations where LMS exceeded 50% of the maximum LMS for that neuron, were used to calculate PD and V, respectively. (E) Quantification of receptive field directionality index (DI). Red arrows indicate positions where LMS exceeded 50% of the maximum LMS (calculated in panel D). Arrow direction indicates LPD, and arrow length is proportional to LMS. DI was calculated by fitting a sine function (red line) to the mean response for each direction (red stars) across these same locations and measuring the normalized difference between the maximum and minimum values of the fit. (F-K) Example receptive fields from TSDNs (F and I) and looming DNs (G, H, J, K), illustrating variation in receptive field size (black line segments indicating height and width), shape (contour line) and directional organization (arrows, direction and length indicate LPD and LMS respectively). Green arrows in (A, F, G) indicate the target trajectories that elicited the strongest response in these example neurons.

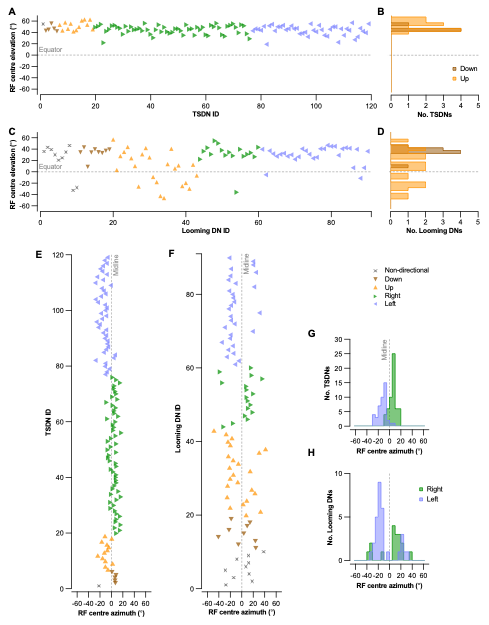

**Figure S3. Receptive field preferred direction varies with position.** Supports Figure 4. (A) Elevation of receptive field centers for individual TSDNs, colored and grouped according to preferred direction. Dashed lines indicate the visual equator. (B) Distributions of receptive field center elevations for upward- and downward-preferring TSDNs. (C and D) Corresponding receptive field center elevations for individual looming DNs (C) and the distributions for upward- and downward-preferring neurons (D). (E and F) Azimuths of receptive field centers for individual TSDNs (E) and looming DNs (F), colored and grouped according to preferred direction. Dashed lines indicate the visual midline. (G and H) Distributions of receptive field center azimuths for rightward- and leftward-preferring TSDNs (G) and looming DNs (H).

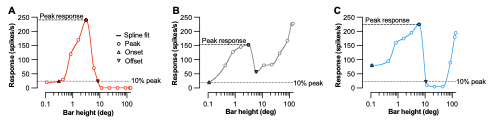

**Figure S4. Quantification of bar size tuning.** Supports Figure 6. Example size tuning curves illustrating the extraction of peak response, onset, and offset thresholds for a TSDN (A), a "TSDN-like" looming DN (B), and a "non-TSDN-like" looming DN (C). Responses at each bar height were fitted with a spline function. Dashed lines indicate the peak response (maximum of the initial peak of the fitted curve) and the 10% threshold used to determine onset and offset. Onset was defined as the bar height at which the fitted response first crossed the 10% threshold, or the smallest bar height tested if the threshold was already exceeded. Offset was defined as the bar height at which the fitted response fell below the 10% threshold after the peak, or the bar height corresponding to the post-peak minimum.
